# Post-traumatic stress symptoms are associated with altered cognitive circuits and threat pathways in chronic pain

**DOI:** 10.64898/2026.02.28.708768

**Authors:** Jennika Veinot, Javeria Ali Hashmi

## Abstract

Chronic pain and post-traumatic stress symptoms (PTSS) frequently co-occur. However, how they interact in the brain to influence cognition and emotion is not well understood. Here, we examined how PTSS affects working memory–related networks during an N-back task. In addition, we examined how these networks interact with subcortical threat regions and influence working memory and pain symptoms. Fifty-three chronic low back pain participants completed the N-back task during fMRI. Brain activation was analyzed in relation to PTSS as both a continuous measure and as high-versus-low groups, using whole-brain parcellation across task loads (FDR corrected). We also examined whether abnormally activated regions were functionally connected to periaqueductal gray subregions, the amygdala, or hippocampus, and how these connections related to PTSS. Although higher PTSS did not affect task performance, it was associated with reduced activation in dorsal and inferior lateral frontal regions during the 3-back condition. PTSS was also associated with increased functional connectivity between the dorsolateral prefrontal cortex and periaqueductal gray, but not with the amygdala or hippocampus. Reduced prefrontal activations and high connectivity with periaqueductal gray predicted higher depression and catastrophizing symptoms. Thus, in chronic pain, PTSS selectively disrupts prefrontal circuits, suggesting that greater trauma symptoms interact with prefrontal circuits when cognitive demand is high. PTSS strengthens coupling between prefrontal regions and brainstem threat/pain circuits, suggesting cognitive-affective coupling. These neural alterations occur even when working memory performance is intact and are linked to higher depression and pain catastrophizing. Larger studies are needed to confirm and clarify these mechanisms.

## Introduction

Chronic primary pain is increasingly recognized as a multidimensional condition that extends beyond persistent sensory symptoms to include alterations in cognition, affect, and brain function [16,88]. Among cognitive domains, working memory is particularly relevant, as it supports attention, goal maintenance, decision-making, and the capacity to engage effectively during treatment [12,62]. Disruption of the affective and regulatory processes that interact with working memory may have important functional consequences for individuals with chronic pain [28,93], contributing to difficulties with emotional regulation, treatment engagement, and vulnerability to mood-related symptoms [12,24,71].

Chronic pain imposes a sustained cognitive load, driven by continuous pain-related signals, high affect and threat monitoring that compete for limited executive resources [24,56,85]. These impairments may emerge primarily under conditions of high task demand [3,24,85]; a relationship that remains to be thoroughly demonstrated. Neuroimaging studies indicate that chronic pain is associated with altered function in frontoparietal networks that support working memory, as well as abnormal interactions with salience and default mode networks [2,51,70,91]. These network-level changes may reflect reduced cognitive efficiency, offering a potential explanation for commonly reported symptoms such as “brain fog,” slowed thinking, and mental fatigue in multiple neuropsychiatric conditions [42,71] but requires systematic investigation. Studying the factors that interact with working memory can help us understand how chronic pain reshapes large-scale brain systems essential for daily functioning.

Post-traumatic stress symptoms (PTSS) are common in people with chronic pain [10,30,80] and may play an important role in contributing to working memory disruption. PTSS is highly prevalent in chronic primary pain and is marked by persistent threat detection, hypervigilance, and intrusive mental content, which increase demands on executive control systems [7,39,44,73,101]. Trauma related alterations in memory function, together with heightened stress system activation, may further impair working memory capacity and cognitive flexibility [14,33,72,79] especially when cognitive load is high [70,71]. Most prior work has focused on studying PTSS and chronic pain as distinct conditions [34], but limited attention was given to PTSS as an exacerbating factor where affect influences cognitive mechanisms to worsen chronic pain symptoms. Examining working memory related brain activations in relation to PTSS may clarify the neural basis of chronic pain and trauma comorbidity.

In this study, we tested working memory function in individuals with chronic pain using an N-back task during functional neuroimaging. We examined whether PTSS modulates task-evoked brain activation during high-demand working memory processing. Furthermore, to test whether interactions between cognitive and affective systems are linked with trauma and pain symptoms and contributes to chronic pain symptoms, we examined the connectivity between cognitive regions that showed aberrant activations during the working memory task and subcortical regions involved in threat and affect processing (i.e. bilateral amygdala, hippocampus, dorsolateral/lateral PAG, ventrolateral PAG). Thus, by assessing PTSS-related abnormalities in brain activation during a working memory task and PTSS-associated alterations in functional connectivity, our aim was to test whether trauma-related processes disrupt cognitive control networks. We hypothesized that higher PTSS would reduce recruitment of working memory regions and enhance the coupling of affected regions with threat-related circuitry.

## Methods and Materials

### Study Data

This study is a component of a larger study directed at developing biopsychosocial and neurological markers for factors associated with treatment failure in chronic back pain (clinicalTrials.gov: RCT #NCT02991625). The main goal of this project is to study the scope and limits of neuroimaging for identifying reproducible and reliable findings from brain data that can pinpoint chronic pain mechanisms. In the present study, we investigate the role that experiencing PTSS has on brain regions involved in working memory in a chronic low back pain population.

### Participants

60 participants (45 women, 15 men) were recruited through advertisements and contact with the Pain Management Unit in Nova Scotia Health, Central Zone (Halifax, NS, Canada). Inclusion criteria for participants were as follows: (1) between 18 and 75 years of age; (2) right-handed; and (3) suffering from chronic low back pain for at least six months with an average reported pain of 4/10 on the brief pain inventory (BPI). Participants were excluded if they had: (1) contraindications for MRI scanning; (2) sensory loss; (3) a history of cardiac, respiratory, or nervous system disease; (4) visual impairment that could not be corrected with lenses; or (5) actively participated in a chronic pain relief treatment program in the last two months. The research protocol was approved by the Nova Scotia Health Authority Research Ethics Board.

### Assessment of Clinical Variables

Clinical variables were assessed through a series of validated self-report questionnaires through REDCap (https://www.project-redcap.org). The current study uses data obtained from the following questionnaires: Brief Trauma Questionnaire (BTQ) [75], PTSD Checklist for DSM-5 (PCL-5) [96], Brief Pain Inventory (BPI) [18], McGill Pain Questionnaire Short-Form (MPQ) [64], State-Trait Anxiety Inventory (STAI) [82], Beck’s Depression Inventory (BDI) [9], Oswestry Low Back Pain Questionnaire [25], and Pain Catastrophizing Scale (PCS) [83].

### Assessment of Traumatic Events and PTSS

Traumatic events were assessed through the Brief Trauma Questionnaire (BTQ) [75]. A tally of the number of participants who answered “yes” for each type of traumatic event and follow-up question outlined in the Brief Trauma Questionnaire can be found in Supplementary Table 1.

PTSS was measured with the PTSD Checklist for DSM-5 (PCL-5). This questionnaire is a 20-item self-report measure that assess the presence and severity of PTSD symptoms, according to the DSM-5 criteria for PTSD [96]. It can be used for quantifying and monitoring symptoms over time, screening individuals for PTSD, and assisting in making a provisional diagnosis of PTSD. The PCL-5 may be used to help determine the appropriate next steps or treatment options for an individual: A total score of 31-33 or higher suggests that a patient may benefit from PTSD treatment, whereas a score below that threshold may indicate that the patient either has subthreshold symptoms of PTSD or does not meet criteria for PTSD [96]. It is important to note that scoring above the cut-point score does not equal an automatic PTSD diagnosis. PTSS can vary in severity and do not always meet full diagnostic criteria for PTSD. It has been acknowledged that assessing PTSS dimensionally, rather than relying solely on categorical PTSD diagnoses, provides a more sensitive and precise measure of symptom severity, captures heterogeneity across individuals, and avoids misclassification due to overlap with other psychiatric disorders [15,55]. As such, here we used a cut-point score of less than 31 to group participants into low post-traumatic stress symptoms (“low PTSS”) and greater than or equal to 31 to group participants into high post-traumatic stress symptoms (“high PTSS”) groups for a between-groups sub analysis.

PTSS in this questionnaire are divided into 4 clusters of symptom type: intrusions/re-experiencing, avoidance, changes in cognition and mood, and arousal/reactivity. Total scores for each of the clusters were calculated for *post hoc* analyses.

### Assessment of Working Memory

#### N-back task

The N-back working memory task is used in assessing working memory ability with fMRI-based neuroimaging [36,45,47,100]. Subjects are presented a sequence of stimuli, one at a time, and are instructed to respond with a button press when the current stimulus is the same as the stimulus presented *n* trials prior. Prior to the start of the experiment, participants were presented with instructions on screen and allowed to practice one block of each task type.

All participants underwent functional MRI scanning while completing a visual N-back letter task that pseudorandomly presented participants with four of each of the following task types: 0-back, 1-back, 2-back, and 3-back. The task was administered across two consecutive scans, each lasting approximately nine minutes, with the 16 task blocks pseudorandomly divided between the two scans. Prior to each block, an instructional screen was presented for 4.75 s indicating which task type was about to occur, and on 0-back trials, it also indicated what the target stimulus letter was (Figure 1). During the task, a letter stimulus (Q, W, R, S, T) was presented in the center of the visual field for 0.475 s followed by a fixation cross for 1.9 s prior the next letter stimulus. Task blocks ranged from 37.5 s (0-back task) to 44.7 s (3-back task) and were followed by an 18 s rest period. The task was designed with fixed proportions of 33% target stimuli and 67% nontarget stimuli. The task was administered using Presentation software (Neurobehavioral Systems). Responses were recorded using an MR-compatible button press. Working memory accuracy scores were calculated from the raw data output for the task and were used as the primary behavioural outcome. Repeated measure analysis of variance (RMANOVA) for accuracy scores was performed across the four task types to assess task difficulty using SPSS (v24; IBM; Armonk, NY, USA), first using PTSS groups as a between-subject factor, and then using PTSS score as a covariate. Other metrics we observed included total omission errors (non-responses to target stimuli), commission errors (responses to nontarget stimuli), total hits (responses to target stimuli), and total correct rejections (non-responses to nontarget stimuli).

**Figure 1.**
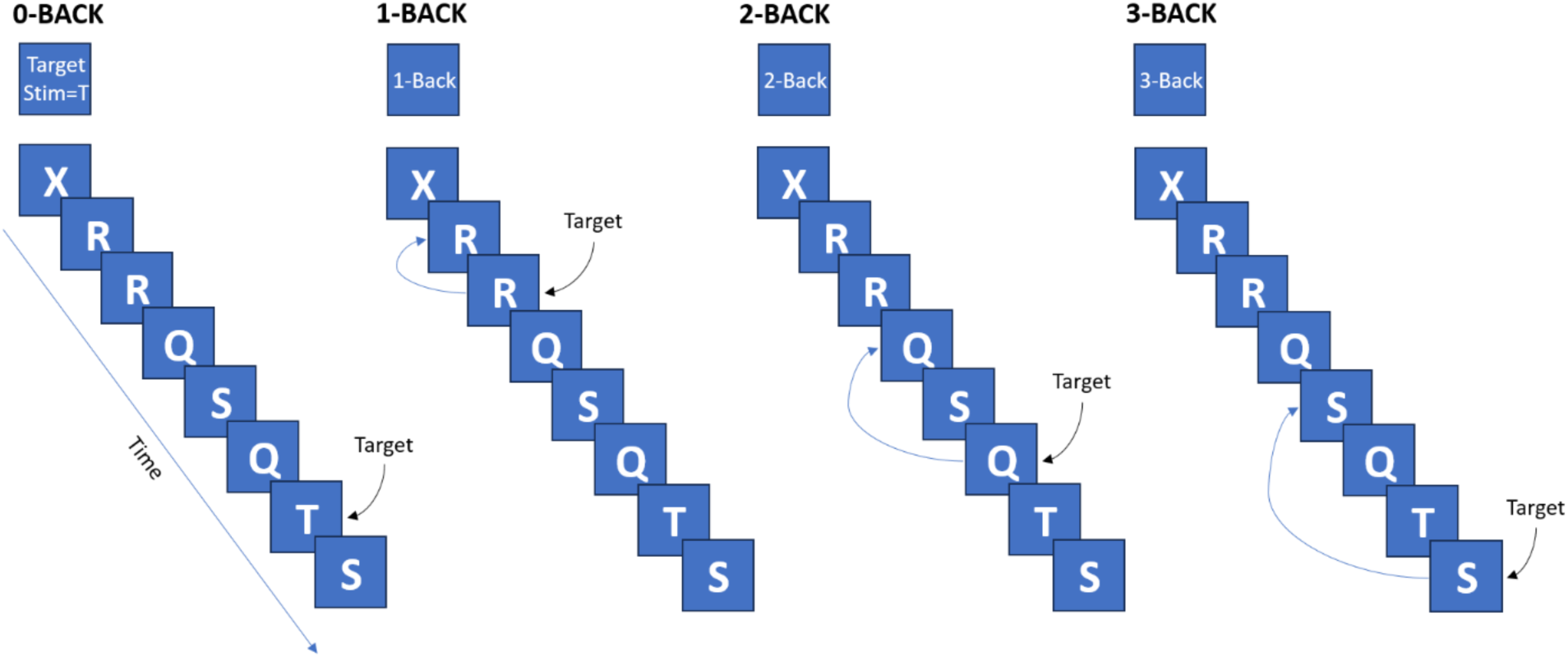
Visual schematic of the four levels of the N-back task paradigm employed in this study.

### fMRI Data Acquisition

Functional data was collected from a 3.0T scanner (Discovery MR750, GE Healthcare; Waukesha, WI, USA) with a 32-channel head coil (MR Instruments, Inc., Minneapolis, MN, USA) at the Biomedical Translational Imaging Centre (BIOTIC) in the Halifax Infirmary, QEII Health Sciences Centre, Halifax, NS, Canada. All acquisition and processing parameters were retained from the parent study [58].

The following parameters were set when acquiring T1-weighted brain images (GE sequence IR-FSPGR): field of view=224×224 mm; in-plane resolution=1mm×1mm; slice thickness=1.0mm; TR/TE=4.4/1.908 milliseconds; flip angle=9°. BOLD (blood oxygenation level dependent) signal sequences for fMRI were acquired using a multi-band EPI sequence as follows: field of view=216×216 mm; in-plane resolution=3mm×3mm; slice thickness=3.0mm; TR/TE=950/30 milliseconds; SENSE factor of 2; acceleration factor of 3. There were 500 total volumes for the resting state scan, 533 volumes for the first task scan and 543 volumes for the second task scan. Reverse phase encoded images were also acquired so FSL’s *topup* could be used for distortion correction.

### fMRI Data Preprocessing

Data were preprocessed with AFNI, FSL and FreeSurfer [20,29,46] using scripts based on those provided by 1000 Functional Connectomes Project [13].

The T1 anatomical image was preprocessed using FreeSurfer’s autorecon1 sequence, which includes motion correction, intensity normalisation, and Talairach transformation. A mask was then generated for stripping the skull away from the image, leaving only the brain; this mask was reoriented to match the original scan then used to crop it. This skull-stripped image was retained for later use.

The functional data were then preprocessed. First, they were corrected for field map-based distortion using topup. Next, several steps were taken using AFNI: (1) discarding the first five EPI volumes to allow for signal equilibration; (2) rigid-body motion correction of time series by aligning each volume to the mean image using Fourier interpolation; (3) skull stripping; and (4) getting an eighth volume for use in registration, as instructed in the scripts provided by the 1000 Functional Connectomes Project. Following that, FSL was used for: (5) spatial smoothing using a Gaussian kernel of full-width half-maximum=6 mm; (6) grand-mean scaling of the voxel value; (7) temporal filtering (0.005-0.3Hz); and (8) removing linear and quadratic trends.

Next, FLIRT (a part of FSL) was used to perform registration of functional images to the Montreal Neurological Institute MNI152 standard template. This included: (1) registration of native-space structural image to the MNI152 2mm template using twelve degree of freedom (df) linear affine transformation; (2) registration of native-space functional image to high-resolution structural image with six df linear transformation; and (3) computation of native-functional-to-standard-structural warps using by concatenating matrices computed in steps (1) and (2).

Finally, several sources of noise were removed in native functional space. Six motion parameters – movement in the x, y, and z planes as well as their rotations, roll, pitch, and yaw – were generated during the motion correction step of the preprocessing. The participant’s T1 image was then segmented into cerebrospinal fluid, grey matter, and white matter, using a tissue-type probability threshold of 80%. From this, the mean signal from the cerebrospinal fluid and white matter, and the mean global signal, were calculated as further nuisance variables. The variables were used in a regression model and the resulting residuals were then registered back into MNI space for time series extraction.

For data quality verification, maximum framewise displacement (FD) and DVARS (difference of volume N to N+1), were calculated. In addition, FSL’s motion outlier detection was used to identify participants with high motion. Participants were to be excluded if they had a maximum FD above 3mm or DVARS outliers detected in more than 10% of the acquired data [74]. No participants were excluded due to motion. Finally, motion was added as a nuisance covariate to verify that the main findings were not due to participant variability in motion.

### Parcellation and Time Series Extraction

For defining regions of interest (ROIs), time series were extracted using a previously reported parcellation scheme optimized for pain studies (Optimized Harvard-Oxford parcellation [37,38]; consisting of 130 bilateral brain regions and the brain stem. Additionally, four periaqueductal gray (PAG) regions (bilateral dorsolateral/lateral and ventrolateral) were defined and extracted based on previously published studies [19,59,95] (Supplementary Figure 1), resulting in 135 ROIs extracted. The BOLD time series were extracted from each voxel within each parcel and averaged. This procedure was done using a BASH script with functions from the FSL library. Given the PAG is subject to motion artifacts, we tested whether the findings from PAG could be replicated by using an ROI covering the entire brainstem as a control, as previously published[95]. We did not check for effects of smoothing on PAG connectivity results since in previous investigations, we found dissociable effects of PAG subregions that were not statistically related to smoothing [1,95].

### N-back Task Evoked Changes in BOLD Activations

To investigate which brain regions were modulated by working memory task load, the portions of BOLD timeseries spanning the four levels of N-back task load were selected and extracted: 0-back, 1-back, 2-back, and 3-back using MATLAB. To account for the hemodynamic response delay of the BOLD signal, the start points of each event were shifted by 6 TRs [57]. The appropriateness of the added delay was confirmed by visualising the time response curves. Activations were quantified by averaging responses over all four trials of each task type.

Whole-brain analyses of paired-samples t-tests contrasting activations during the 3-back task against activations during the lower task loads (2-back, 1-back and 0-back) were performed across all participants using MATLAB to see which ROIs were modulated by each increase in working memory load. All results were FDR corrected at *q*=0.05. Wilcoxon rank sum test was used between PTSS groups in MATLAB to observe if any patterns of brain activations during each separate task load were hindered by presence of PTSS. Regions that differed between groups following FDR correction were further analyzed using repeated measures ANOVAs (including using PTSS groups as a between-subject factor) and were selected to observe task-state functional connectivity between threat circuitry regions.

### ROI-based Task-State Functional Connectivity

To determine functional connectivity during task scans, a zero-lag Pearson correlation matrix was calculated using MATLAB for the entire task between the mean BOLD signals that were determined to be modulated by task load and impacted by PTSS (in the BOLD activations analysis), and ROIs involved in threat circuitry (bilateral amygdala, hippocampus, dorsolateral/lateral PAG, ventrolateral PAG, and as a control, the entire brainstem). Results were FDR corrected at *q*=0.05. Functional connectivity values between regions that survived FDR correction were correlated against PTSS as well as patterns of brain activations during the 3-back task to observe if the FC values predicted PTSS and/or patterns of brain activity modulated by PTSS using SPSS. Finally, *post hoc* correlations were conducted between the FC values and PTSS symptom subcategories that were found to be significantly correlated after FDR correction.

## Results

### Clinical and behavioral results

Data was collected from 60 participants (45 women, 15 men) with chronic low back pain. One participant dropped out due to claustrophobia, leaving 59 participants who completed the study. Of the remaining participants, five participants could not complete the *N*-back task in the MRI due to equipment failure, so they completed it outside of the scanner, and one participant’s MRI data was not used due to an undisclosed previous brain resection, resulting in MRI task data from 53 participants. Data from these participants were analysed based on groupings by PTSS severity (High PTSS *N* =14, Low PTSS *N* =39) and by using PTSS as a continuous variable.

Higher PTSS scores in this data were linked with high affective load (Table 1). Specifically, PTSS observed as a continuous variable correlated significantly with trauma exposure (BTQ: *r*=0.612, *p*<0.001), depression scores (BDI: *r*=0.709, *p*=<0.001), state anxiety (STAI-S: *r*=0.612, *p*<0.001), trait anxiety (STAI-T:*r*=0.700, *p*<0.001), pain catastrophizing (PCS: *r*=0.618, *p*<0.001), affective pain (MPQ-Affect: *r*=0.417, *p*<0.001), but not with sensory pain (MPQ-sensory: *r*=0.137, *p*>0.05), chronic pain intensity (MPQ-VAS: *r*=-0.048, *p*>0.05), or disability (Oswestry: *r*=-0.273, *p*>0.05). When comparing between groups, the high PTSS group had significantly greater exposure to trauma (BTQ: *p*<0.001), state anxiety (STAI-S: *p*<0.001), trait anxiety (STAI-T: *p*<0.001), depression (BDI: *p*<0.001), pain catastrophizing (PCS: *p*<0.001), and affective pain (MPQ-Affect: *p*=0.01) compared to the low PTSS group (see Table 1). In addition, among traumatic events experienced across participants (Supplementary Table 1), the most reported was sexual assault, which was reported in 59% of the sample.

**Table 1.** Demographic and clinical metrics across all participants and across PTSS groups. * = *p* < 0.05.

|  | *All*<br>*Participants* | *High PTSS* | *Low PTSS* |
| --- | --- | --- | --- |
| **N** | 59 | 16 | 43 |
| **% Women** | 75% | 81.3% | 72.1% |
| **Age** | 41.2 ± 12.2 | 35.2 ± 10.4* | 42.8 ± 12.1* |
| **Exposure to Trauma (BTQ)** | 3.6 ± 3.1 | 6.0 ± 2.6* | 2.7 ± 2.8* |
| **Post-traumatic Stress**<br>**Symptoms** | 25.3 ± 14.0 | 44.5 ± 9.0* | 18.2 ± 7.1* |
| **State Anxiety** | 46.4 ± 10.8 | 56.6 ± 7.1* | 42.5 ± 9.4* |
| **Trait Anxiety** | 45.1 ± 12.8 | 40.6 ± 11.5* | 57.0 ± 7.3* |
| **Depression** | 16.2 ± 10.3 | 26.9 ± 9.1* | 12.1 ± 7.4* |
| **Disability** | 15.7 ± 7.5 | 17.3 ± 5.6 | 15.1 ± 1.3 |
| **Pain Catastrophizing** | 22.2 ± 10.5 | 32.0 ± 9.5* | 18.4 ± 8.2* |
| **McGill Pain Affective** | 5.22 ± 3.07 | 6.88 ± 2.87* | 4.60 ± 2.93* |
| **McGill Pain Sensory** | 16.19 ± 6.01 | 16.81 ± 4.65 | 15.95 ± 6.49 |
| **McGill Pain Total** | 21.41 ± 8.20 | 23.69 ± 6.06 | 20.55 ± 8.79 |
| **BPI Average Chronic Pain**<br>**Intensity** | 53.1 ± 14.3 | 52.31 ± 15.1 | 53.33 ± 14.3 |
| **Self-Identified Race** |  |  |  |
| *Black* | 2 | 0 | 2 |
| *First Nations* | 2 | 1 | 1 |
| *Latin* | 1 | 1 | 0 |
| *Metis* | 4 | 1 | 3 |
| *South Asian* | 2 | 0 | 1 |
| *White/European* | 49 | 13 | 36 |

### N-back accuracy decreases with task load

Repeated measures ANOVA determined that N-back accuracy differed significantly across the four task types (F_(2.447,141.904)_ = 65.834, *p* <.001; Figure 2). Post hoc pairwise comparisons using Bonferroni correction showed decreased N-back accuracy with each increasing task type at *p*<.001, except for the pairwise comparison between the 0-back and 1-back tasks which was significant at *p*=0.002. Across all participants, mean overall accuracy for the N-back task was 0.922 (SD = 0.0439). Notably, there were no differences in accuracy or any other measured working memory metric between PTSS groups (Table 2). Further, no linear relationships existed between any working memory metric and total PTSS in this sample (*p*>0.05). Finally, change in accuracy with task load was not associated with PTSS scores either as a covariate (p>0.05) or between groups (p>0.05) when assessed with a repeated measures ANOVA.

**Figure 2.**
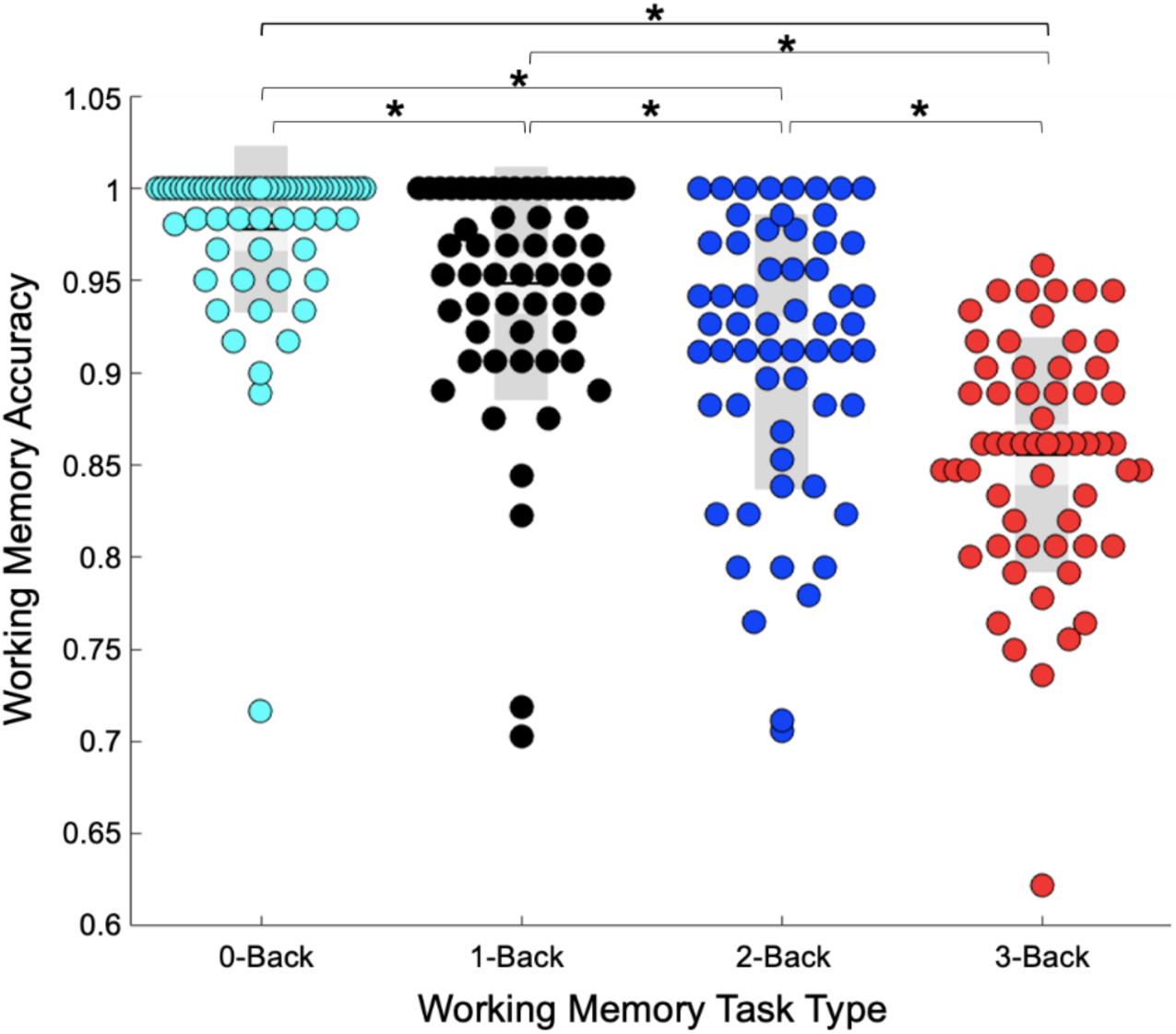
Working memory accuracy scores for 0-back task (turquoise), 1-back task (black), 2-back task (blue) and 3-back task (red). * = *p*<.05.

**Table 2.** N-back Task metrics for all participants and for PTSS groups.

|  | *All*<br>*Participants* | *High PTSS* | *Low PTSS* |
| --- | --- | --- | --- |
| ***Overall Accuracy*** | 0.922 ± 0.0439 | 0.9274 ±<br>0.0363 | 0.9196 ±<br>0.0467 |
| ***0-Back Accuracy*** | 0.978 ± 0.0448 | 0.9537 ±<br>0.0768 | 0.9864 ±<br>0.0228 |
| ***1-Back Accuracy*** | 0.948 ± 0.0629 | 0.9688 ±<br>0.0378 | 0.9411 ±<br>0.0692 |
| ***2-Back Accuracy*** | 0.911 ± 0.0742 | 0.9392 ±<br>0.5505 | 0.8997 ±<br>0.0778 |
| ***3-back Accuracy*** | 0.855 ± 0.0635 | 0.8607 ±<br>0.0599 | 0.8534 ±<br>0.0659 |
| **Hits** | 0.8185 ±<br>0.1276 | 0.8481 ±<br>0.0723 | 0.8066 ±<br>0.1420 |
| **Misses** | 0.1815 ±<br>0.1276 | 0.1519 ±<br>0.0723 | 0.1934 ±<br>0.1420 |
| **Correct Rejections** | 0.9653 ±<br>0.0232 | 0.9620 ±<br>0.0293 | 0.9665 ±<br>0.0214 |
| **False Alarms** | 0.0340 ±<br>0.0235 | 0.3804 ±<br>0.2926 | 0.0326 ±<br>0.0293 |

### Engagement of working memory networks across N-back loads

First, we examined if increasing task load engaged the working memory network, by identifying regions showing significant load-dependent activation across the whole brain in the entire group (N=53, FDR corrected *q*=0.05, Figure 3). The comparisons showed that the 3-back task load activated 30 regions significantly more relative to 0-back task load (Table 3) affecting the frontal parietal, dorsal attention, cingulate and reward processing regions. Similarly, the 3-back task load activated 19 regions significantly more relative to the 1-back, activating the dlPFC, cingulate and visual and posterior default mode regions. Finally, the 3-back versus 2-back contrast revealed only an occipital region significantly more active in the 3-back (Table 3). As shown in Figure 3, these regions included frontal, parietal, and visual cortices, consistent with the increased cognitive demands of the 3-back condition. The regions identified in the 3-back versus 0-back contrast showed substantial correspondence with established working memory networks, with approximately 70% overlap with voxels from NeuroSynth’s Working Memory Association Map, whereas the 3-back versus 1-back contrast demonstrated more limited correspondence, with 26% overlap. The single occipital region identified in the 3-back versus 2-back contrast also overlapped with the map. Notably, the highest correspondence was observed for the 3-back condition, indicating that this contrast best captures canonical voxel-wise activation patterns of working memory and supporting its use for subsequent analyses.

**Figure 3.**
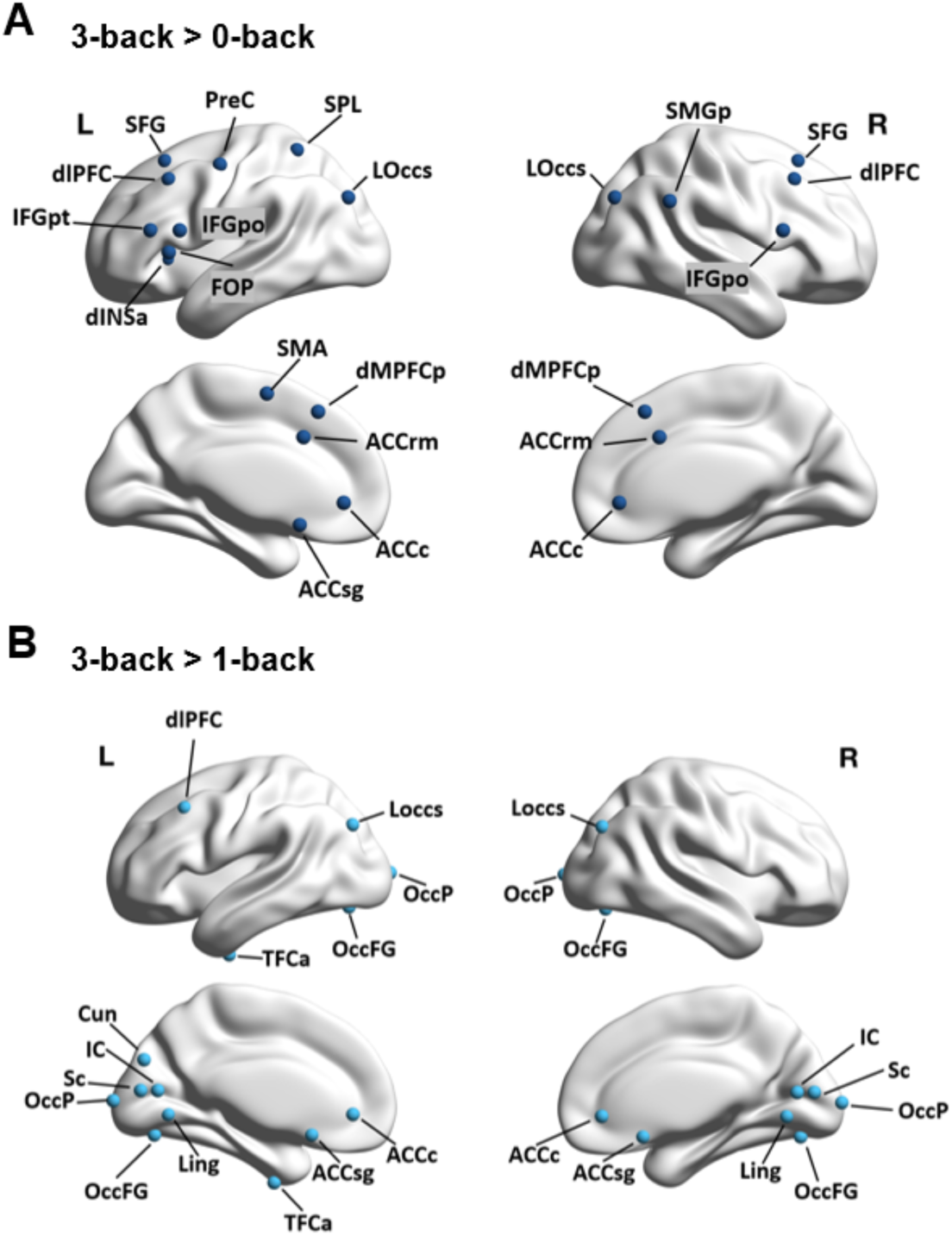
**[A]** Brain regions that were more active during the 3-back task compared to the 0-back task following whole-brain FDR correction. **[B]** Brain regions that were more active during the 3-back task compared to the 1-back task following whole-brain FDR correction. *dlPFC=dorsolateral prefrontal cortex, dMPFCp=posterior dorsomedial prefrontal cortex, dINSa=dorsal anterior insula, ACCc=caudal anterior cingulate cortex, ACCsg=subgenual anterior cingulate cortex, ACCrm=rostromedial anterior cingulate cortex, TFCa=anterior temporal fusiform cortex, Ling=lingual gyrus, IC=intracalcarine cortex, Loccs=superior lateral occipital cortex, Cun=cuneal cortex, Sc=supracalcarine cortex, SFG=superior frontal gyrus, SPL=superior parietal lobule, SMGp=posterior supramarginal gyrus, PreC=precentral gyrus, IFGpt=inferior frontal gyrus pars triangularis, IFGpo=inferior frontal gyrus pars opercularis, FOP=frontal operculum, OccP=occipital pole, OccFG=occipital fusiform gyrus, L=left hemisphere, R= right hemisphere*.

**Table 3.**
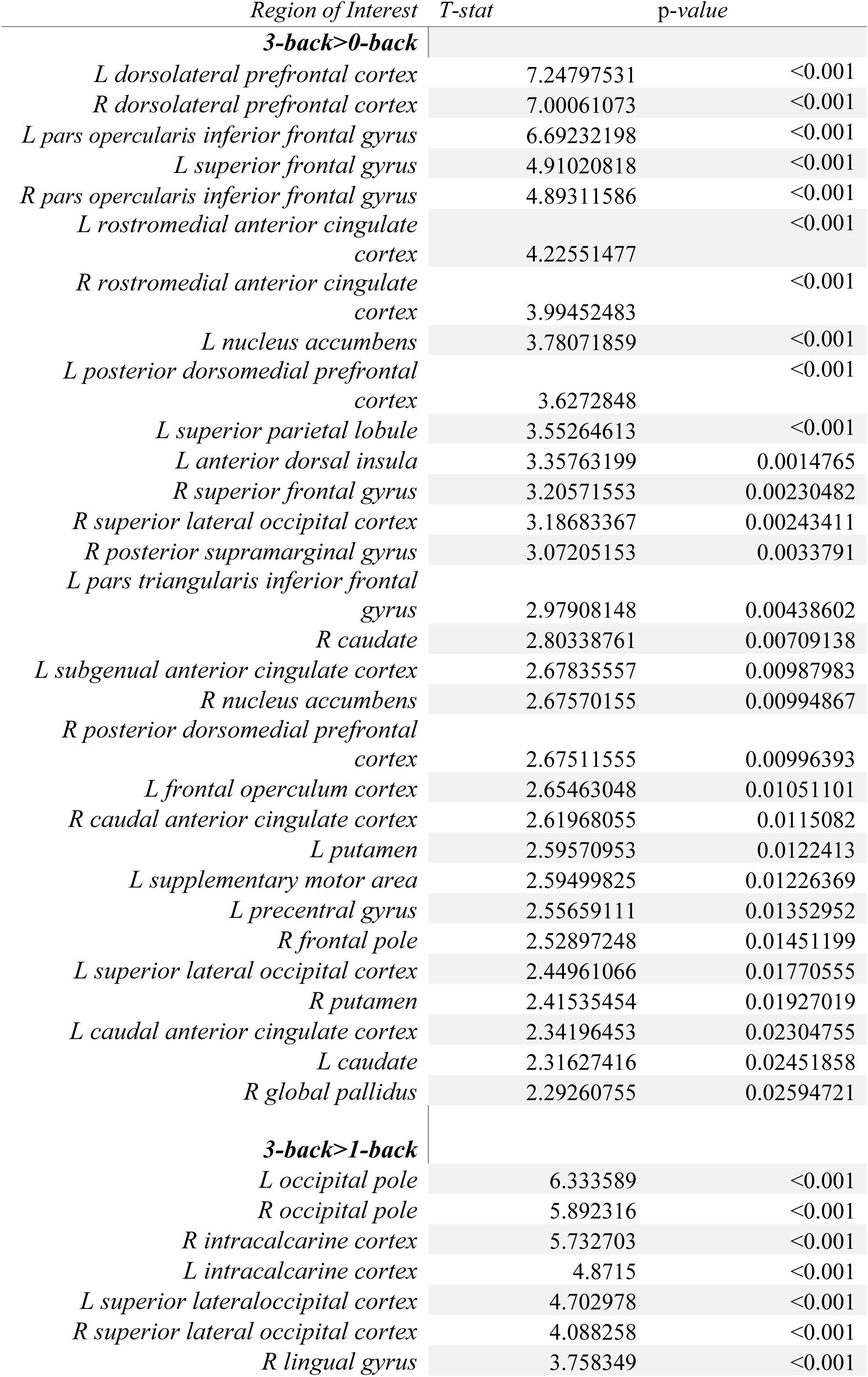
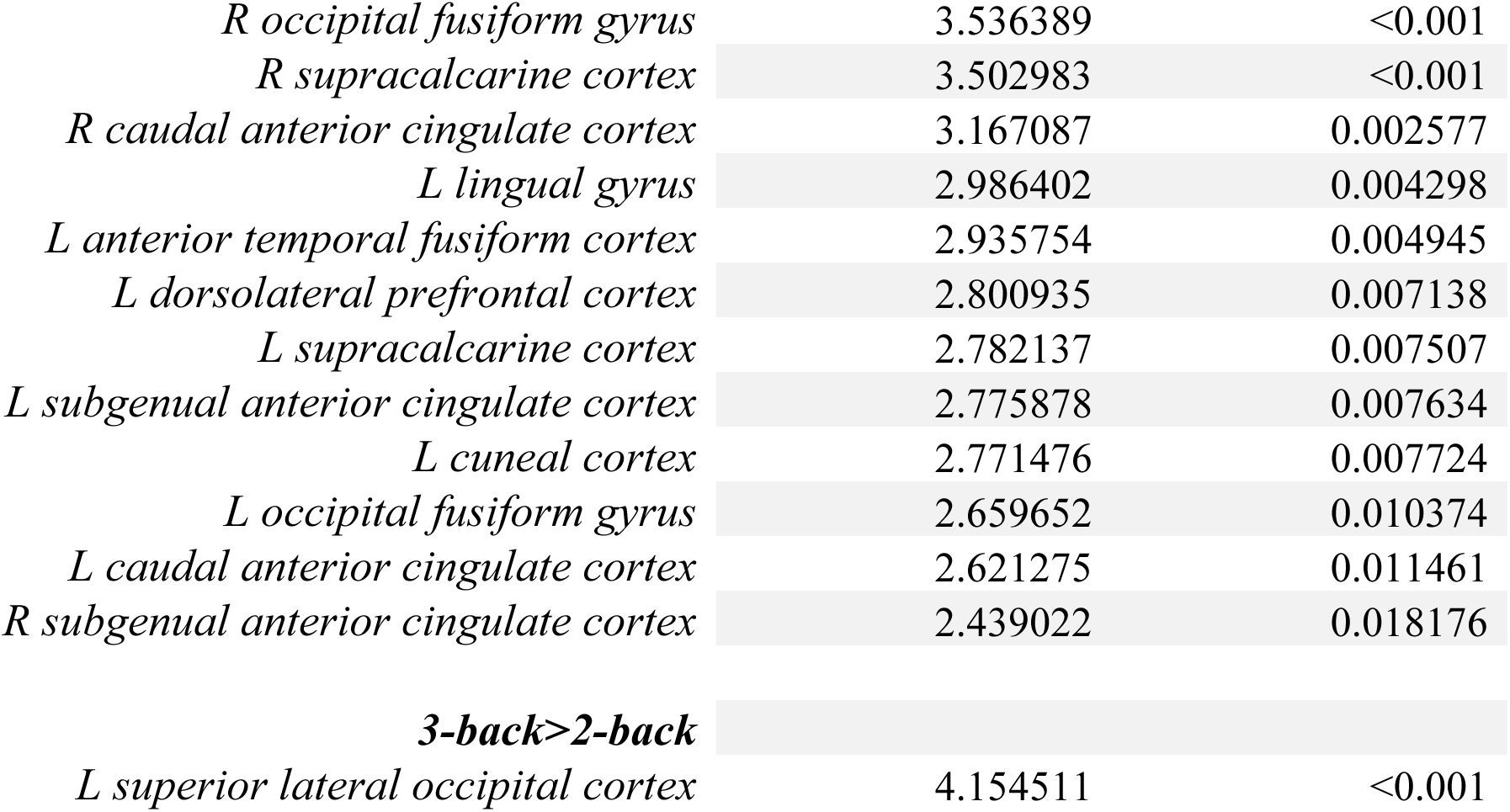
Brain regions that were more active during the 3-back task compared to the 0-back, 1-back and 2-back tasks following whole-brain FDR correction.

### PTSS modulates prefrontal and parietal activation during high cognitive demand

To address PTSS-related effects on working memory networks, we examined PTSS group differences separately at each task load using whole-brain FDR-corrected analyses (q<0.1), rather than restricting inference to regions defined by task-load contrasts. This approach avoided allowed us to confirm whether PTSS-related effects were load-specific or present across the different levels of cognitive demand. Given the modest sample size, we focused interpretation on effects that were reproducible across both group-based and continuous PTSS analyses. As a secondary step, we verified whether the regions that showed groupwise differences in activation showed a task load dependent effect when tested as a repeated measures ANOVA.

Whole-brain FDR-corrected group comparisons revealed significant PTSS-related differences in activations only during the 3-back condition (Figure 4, whole brain FDR corrected, q<0.1) and limited to three regions: the left dorsolateral prefrontal cortex (dlPFC; *z* = 3.12, *p* = 0.0018), the left angular gyrus (*z* = 3.08, *p* = 0.0021), and the left pars opercularis of the inferior frontal gyrus (IFGpo; *z* = 3.34, *p* < 0.001), with reduced activation in the high PTSS group in all three regions (see Figure 4A,B,D,E,G,H).

**Figure 4.**
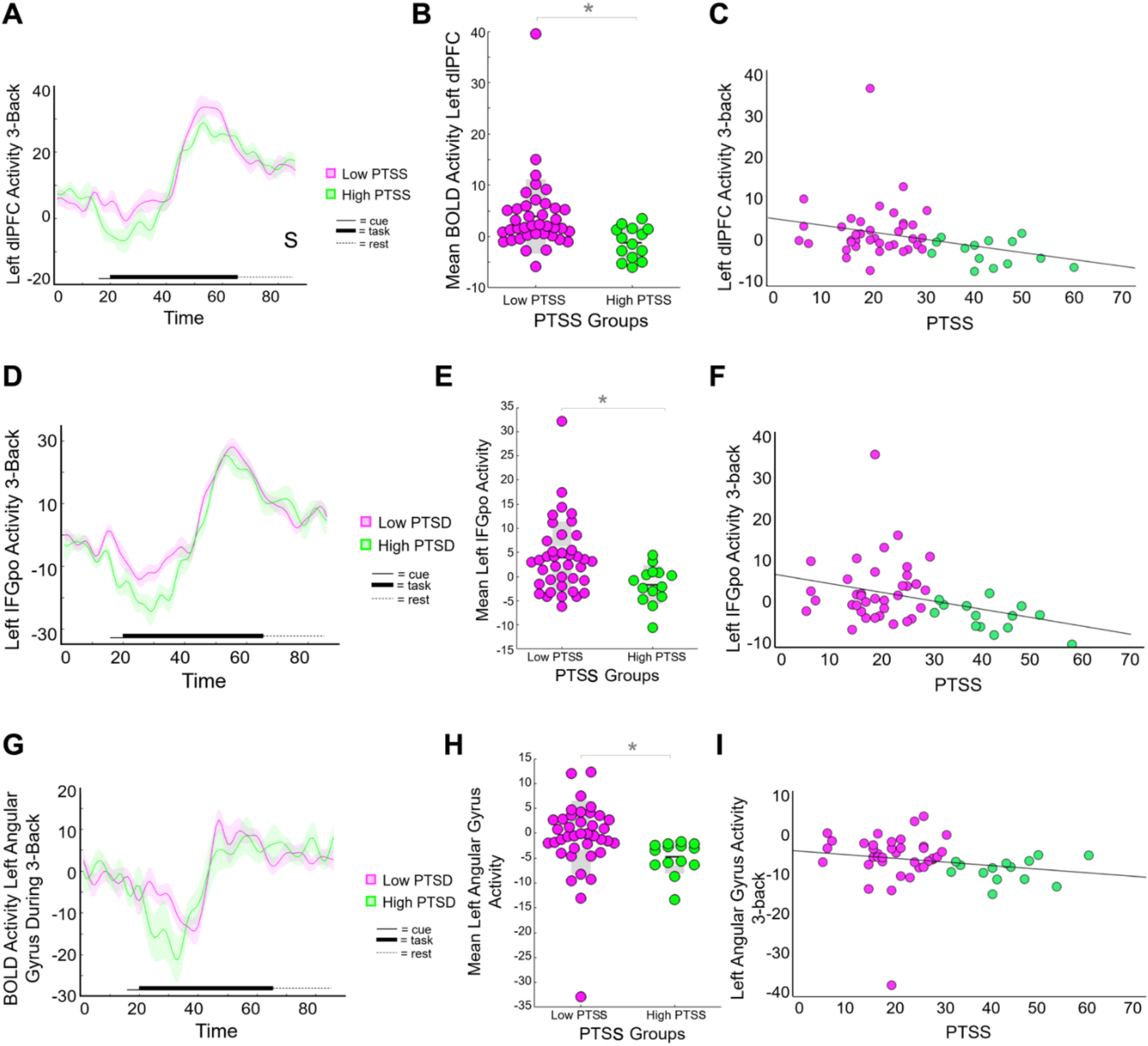
**[A]** Averaged BOLD activation in the left dlPFC across the entire task durations for low PTSS (pink) and high PTSS (green). **[B]** Individual BOLD activations of the left dlPFC during the 3-back for the low PTSS group (pink) and the high PTSS group (green). **[C]** Individual averaged BOLD activity in left dlPFC during 3-back task by severity of PTSS. **[D]** Averaged BOLD activation in the left IFG across the entire task durations for low PTSS (pink) and high PTSS (green). **[E]** Individual BOLD activations of the left IFG during the 3-back for the low PTSS group (pink) and the high PTSS group (green). **[F]** Individual averaged BOLD activity in left IFG during 3-back task by severity of PTSS symptoms. **[G]** Averaged BOLD activation in the left angular gyrus across the entire task durations for low PTSS (pink) and high PTSS (green). **[H]** Individual BOLD activations of the left angular gyrus during the 3-back for the low PTSS group (pink) and the high PTSS group (green). **[I]** Individual averaged BOLD activity in left angular gyrus during 3-back task by severity of PTSS. **\*=***p*<0.05*. Pink= low PTSS, green= high PTSS*.

To confirm that the association is not driven solely by grouping, we next tested associations between PTSS and BOLD activations of these regions during each task load. We found that PTSS severity was negatively correlated with left dlPFC activation during the 3-back task (*r*=-0.342, *p*=0.013; Fig. 4C), confirming that the association has a graded relationship. Similarly, PTSS severity was also negatively associated with left inferior frontal gyrus activation during the 3-back (*r*=-0.320, *p*=0.022; Figure 4F), as well as left angular gyrus activation during the 3-back (*r*=-0.348, *p*=0.012; Figure 4I). These associations remained significant after including motion as a covariate (*p*<0.05). PTSS severity assessed as a continuous variable was not associated with activations during 0, 1, or 2-back task loads (*p*>0.05).

### PTSS modulates prefrontal activations when task load is high

Next, to confirm whether the identified regions showed load-dependent group differences, we included all task loads in a single model and tested the task load × group interaction and also visualized the load dependent activations for all participants (Figure 5). Repeated-measures ANOVAs including all four task loads as a within-subject factor and PTSS group as a between-subject factor revealed significant task load × group interactions for the left dlPFC (F_(1.939, 98.898)=_3.60, *p*=0.033) and left IFGpo (F_(2.391, 119.527)_=3.65, *p*=0.014). Notably, no significant task × group interaction was observed for the angular gyrus (*p*>0.05). In addition, the angular gyrus did not show load dependent changes, indicating that the PTSS effects on angular gurus were not related specifically to task load. The group differences remained significantly different for these regions after including motion as a covariate (*p*<0.05). For the dlPFC, post hoc paired t-tests indicated significantly greater activation during the 3-back task (M=2.56, SD=6.74) compared to the 0-back (M=-8.20, SD=6.53; t_(52)_=7.25, *p*<0.001), 1-back (M=-0.30, SD=4.67; t_(52)_=2.80, *p*=0.007), and 2-back (M=0.67, SD=7.12; t_(52)_=2.41, *p*=0.020) conditions (Figure 5A and 5B). Similarly, for the IFGpo, post hoc paired t-tests indicated significantly greater activation during the 3-back task (M=1.47, SD=6.98) compared to the 0-back (M= -8.75, SD=7.19; t_(52)_=6.53, *p*<0.001), and 2-back (M=-0.768, SD=5.95; t_(52)_=2.10, *p*=0.41), but not the 1-back (M=0.234, SD=4.88; t_(52)_=0.992, *p*>0.05) conditions (Figure 5C and 5D). In contrast, left angular gyrus showed no load-dependent modulation (*p* > 0.05), indicating that task load effects were specific to dlPFC and IFGpo (Figure 5E and 5F). Note that negative mean activations do not reflect deactivations during task, but rather a delayed rise in activation relative to task onset, which can result in negative mean values that reflect the temporal dynamics of the hemodynamic response when collapsed across the full-time window.

**Figure 5.**
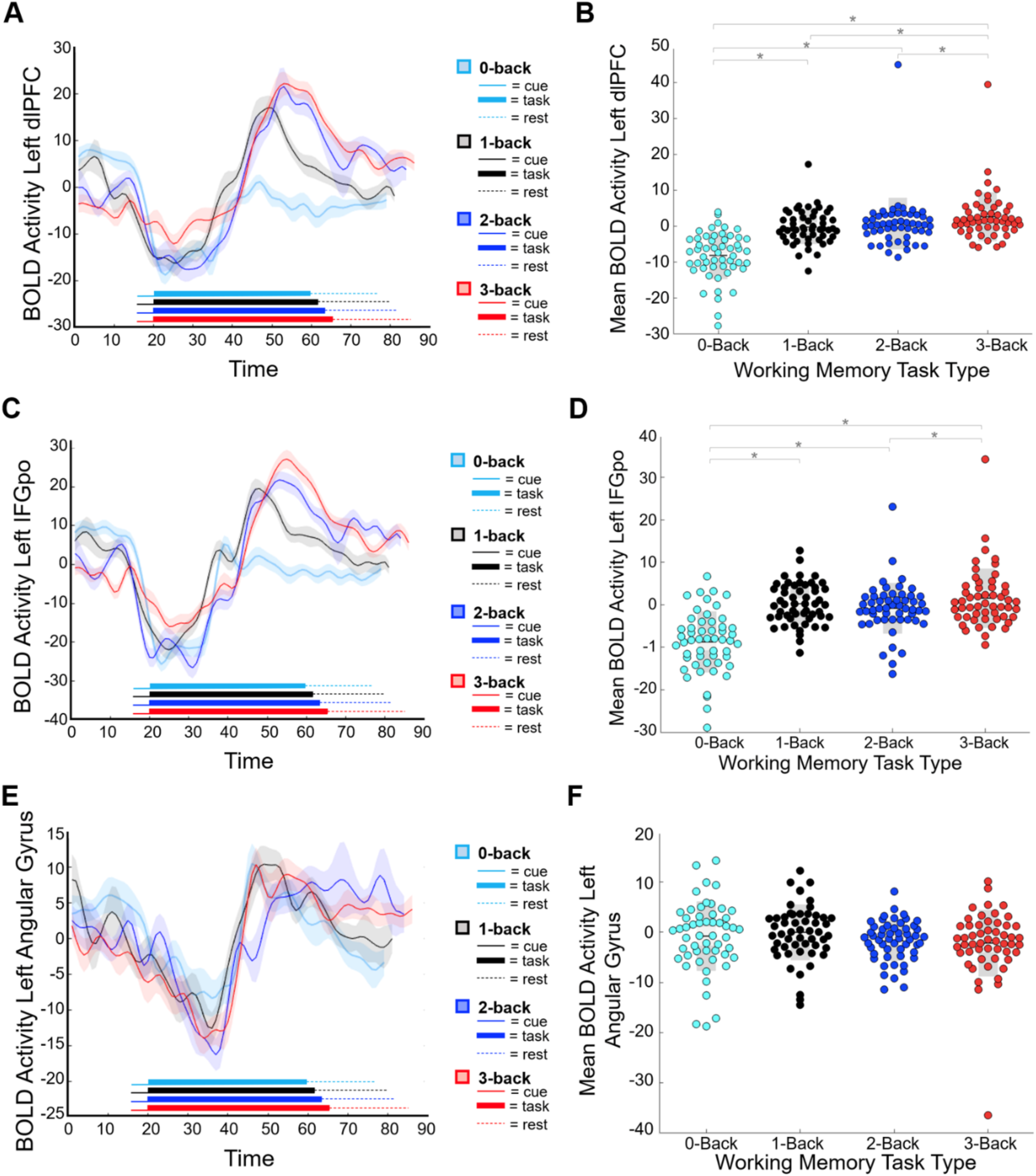
**[A]** Averaged BOLD activation in the left dlPFC across the entire task durations. **[B]** Individual BOLD activations of the left dlPFC for each task type. **[C]** Averaged BOLD activation in the left IFG across the entire task durations. **[D]** Individual BOLD activations of the left IFG for each task type. **[E]** Averaged BOLD activation in the left angular gyrus across the entire task durations. **[F]** Individual BOLD activations of the left angular gyrus for each task type. (0-back=light blue, 1-back=black, 2-back=dark blue, 3-back=red). *=*p*<.05.

We further confirmed that PTSS-modulated regions tracked task load independently of group categorization by repeating the analyses using continuous PTSS scores. These ANOVAs showed a significant main effect of task for the left dlPFC (F_(1.915, 99.583)_=31.592, *p*<0.001) and IFGpo (F_(2.356, 120.134)_=25.324, *p*<0.001), with activation increasing at higher loads, particularly during 3-back, and remaining significant after controlling for motion.

### PTSS modulates functional coupling between cognitive control regions and threat-processing areas

We next evaluated whether regions modulated by PTSS at high task load (left dlPFC, left IFGpo, left angular gyrus) exhibited functional connectivity with threat-processing structures (amygdala, hippocampus, dlPAG, vlPAG), using the whole brainstem as a control. After correction for multiple comparisons, PTSS severity was positively correlated with functional connectivity between left dlPFC – left dl/lPAG (*r*_(51)_= 0.391, *p*=0.004; Figure 6A), as well as left dlPFC – left vlPAG (*r*_(51)_= 0.322, *p*=0.04; Figure 6B), but not for any other functional connections (*p*>0.05). These results remained significant after controlling for motion (*p*<0.05). When these functional connections were compared between PTSS groups (FDR corrected), the high PTSS group showed significantly greater left left dlPFC – left dl/lPAG FC compared to the low PTSS group (low PTSS=-0.0161 ± 0.134, high PTSS=0.0875 ± 0.0927, *p*=0.012; Figure 6C). Similarly, left dlPFC – left vlPAG FC was significantly greater in the high PTSS group (low PTSS = -0.0293 ± 0.115, high PTSS = 0.0684 ± 0.107, *p*=0.032; Figure 6D). No significant group differences emerged for dlPFC connectivity with the left or right amygdala, hippocampus, the right dl/lPAG, right vlPAG, or the control brain stem region (*p*>0.05) Additionally, following FDR-correction, no significant between group differences were found for functional connectivity between threat processing regions and the IFG or angular gyrus (*p*>0.05).

**Figure 6.**
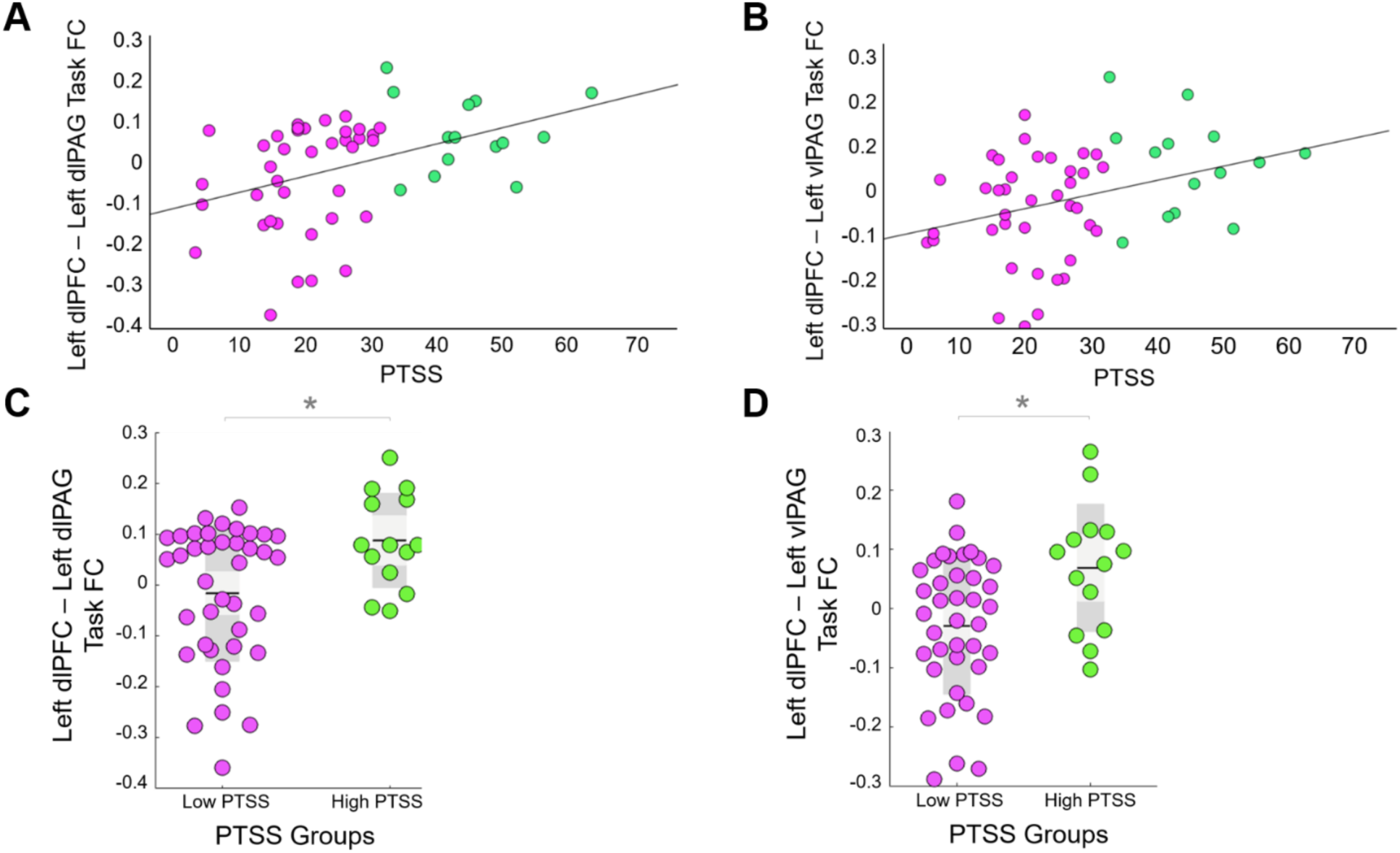
**[A]** Individual averaged task-based functional connectivity between the left dlPFC and left dl/lPAG during the N-back task by severity of PTSS. **[B]** Individual averaged task-based functional connectivity between the left dlPFC and left vlPAG during the N-back task by severity of PTSS. **[C]** Individual averaged task-based functional connectivity between the left dlPFC and left dl/lPAG during the N-back task for the low PTSS group (pink) and the high PTSS group (green). **[D]** Individual averaged task-based functional connectivity between the left dlPFC and left vlPAG during the N-back task for the low PTSS group (pink) and the high PTSS group (green). . *Pink= low PTSS, green= high PTSS*.

Further investigation revealed that low dlPFC activation during the 3-back task was predicted by greater left dlPFC – left dl/lPAG functional connectivity (*r*_(52)_=-0.308, *p*=0.025; Figure 7A), as well as by left dlPFC – left vlPAG functional connectivity (*r*_(52)_=-0.355, *p*=0.009; Figure 7B). Neither left IFG activation nor left angular gyrus activation during the 3-back task was predicted by their connectivity with either PAG subregion (*p*>0.05).

**Figure 7.**
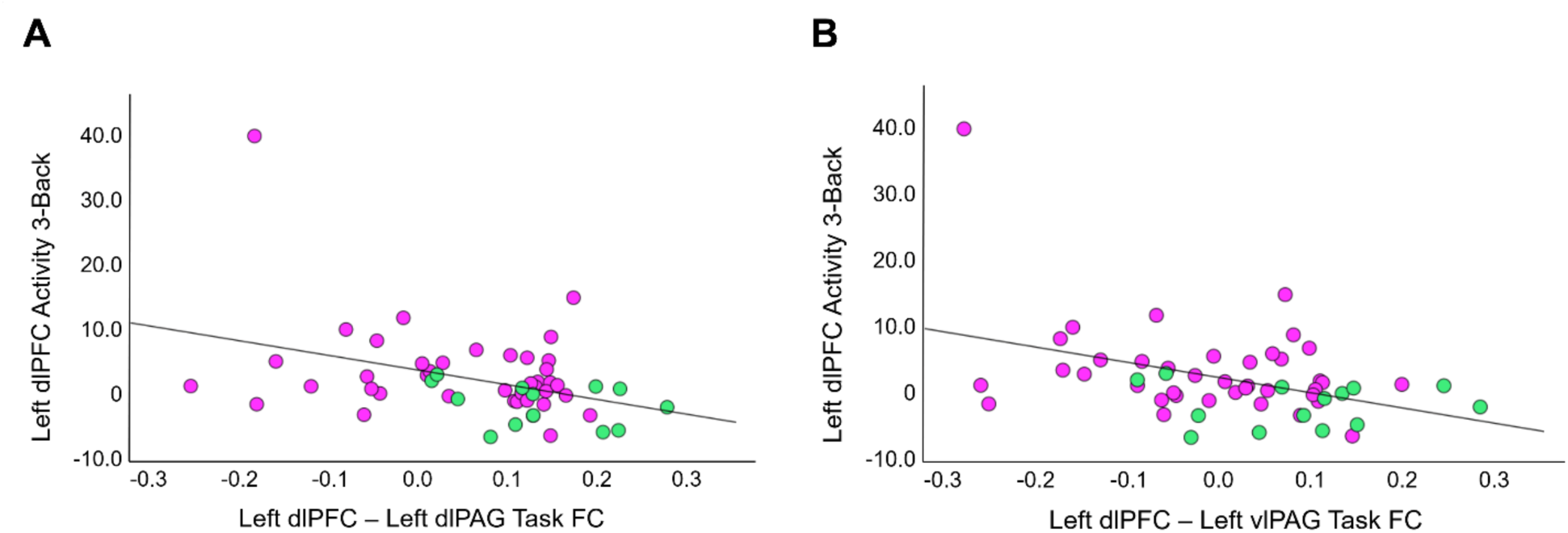
**[A]** Individual averaged left dlPFC activity during the 3-back task by individual averaged task-based functional connectivity between the left dlPFC and left dl/lPAG during the entire n-back task. **[B]** Individual averaged left dlPFC activity during the 3-back task by individual averaged task-based functional connectivity between the left dlPFC and left vlPAG during the entire n-back task. *Pink= low PTSS, green= high PTSS*.

### Behavioral relevance of PTSS-related activation and connectivity

Activations in any of the identified brain regions shown to be affected by high PTSS were not associated with total working task accuracy (*p*>0.05) suggesting that reduced activations did not affect task performance. Moreover, these activations also did not predict chronic pain intensity (MPQ-VAS, p>0.05) or disability (Oswestry, *p*>0.05). In comparison, activations in dlPFC during the 3-back condition was associated with higher PTSS (*r*=-0.342, *p*=0.013), increased pain catastrophizing (r= -0.372, *p*=0.007) and greater depression symptoms (r= -0.377, *p*=0.006) indicating that the reduced dlPFC activations observed in high PTSS were linked behaviorally with higher affective disorder.

The identified functional connectivity patterns associated with high PTSS also showed an association with affective symptoms. The higher left dlPFC – left dl/lPAG (*r*= -0.309, *p*=0.026) and left dlPFC – left vlPAG (*r*= -0.280, *p*=0.045) functional connectivity observed in relation with high PTSS were significantly associated with higher depression scores. Pain intensity and anxiety did not show a significant correlation (*p*>0.05).

### dlPFC–PAG subregion connectivity tracks distinct PTSS symptom clusters

We next wanted to investigate whether dlPFC – PAG subregion connectivity correlated differently with specific PTSS subcategories ([A] avoidance, [B] intrusion, [C] arousal and reactivity, and [D] negative changes in cognition and mood). *Post hoc a*nalyses revealed that after multiple comparison correction, left dlPFC – left dl/lPAG FC predicted arousal and reactivity symptoms (*r*_(51)_=0.421, *p*=0.008) and changes in cognition and mood symptoms (*r*_(51)_=0.348, *p*=0.044), but not avoidance symptoms (*r*_(51)_=0.312, *p*=0.096) or intrusion symptoms (*r*_(51)_=0.257, *p*=0.264). Conversely, left dlPFC – left vlPAG FC predicted avoidance symptoms (*r*_(51)_=0.366, *p*=0.032) and changes in cognition in mood symptoms (*r*_(51)_=0.352, *p*=0.044), but not arousal and reactivity symptoms (*r*_(51)_=0.251, *p*=0.288) or intrusion symptoms (*r*_(51)_=0.156, *p*=1.00). Finally, to ensure that dlPFC–PAG subregion connectivity associations with PTSS subcategories were not actually driven by dlPFC task activation, we included dlPFC task activation as a covariate in the analysis. After controlling for left dlPFC activity during the 3-back task and correcting for multiple comparisons, the only correlations that survived were between left dlPFC – left dl/lPAG FC and arousal and reactivity symptoms (*r*_(49)_=0.345, *p*=0.013; Figure 8A), and left dlPFC – left vlPAG FC and avoidance symptoms (*r*_(49)_=0.308, *p*=0.028; Figure 8B).

**Figure 8.**
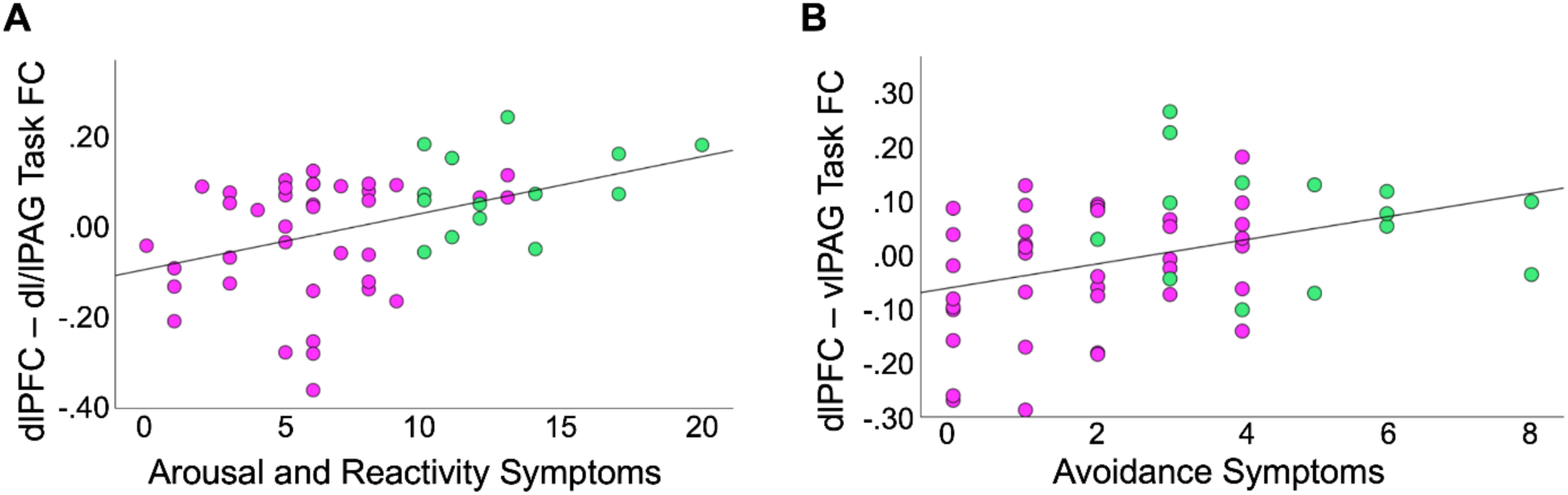
**[A]** Individual averaged task-based functional connectivity between left dlPFC and left dl/lPAG during n-back task by PCL arousal and reactivity symptoms. **[B]** Individual averaged task-based functional connectivity between left dlPFC and left vlPAG during n-back task by PCL arousal and reactivity symptoms.

## Discussion

Our findings indicate that PTSS in chronic low back pain is associated with reduced activation in canonical working memory regions, including the dlPFC and IFGpo, but these differences were evident only under the highest cognitive load. Further, higher PTSS was linked with increased functional connectivity between the dlPFC and the PAG, a key threat-processing region, indicating that individuals with higher PTSS also show heightened engagement with threat circuitry. Importantly, these neural patterns predicted pain affect, depressive symptoms, and pain catastrophizing, but were not associated with working memory performance, chronic pain intensity, or disability. Together, these results suggest that in chronic pain, PTSS involves aberrant prefrontal–threat circuit interactions that are associated with the affective pain dimension.

The dlPFC and IFGpo are central components of lateral prefrontal control networks that support working memory and provide top-down regulation of both emotional reactivity [11,98,103] and descending pain modulation through their interactions with limbic regions and midbrain structures [61,78,84,93]. Substantial evidence shows that these regions shape working memory under increasing cognitive load [6,48], regulate threat-related affect [43,54,65,68], and engage endogenous analgesic mechanisms [19,21,67,86]. Further, their structure and function are consistently altered in chronic pain [41,69,78,102]. In this context, the load-dependent reduction in dlPFC and IFGpo recruitment observed in individuals with higher PTSS points to a selective reduction in regulatory efficiency within prefrontal control systems, but is not a primary limitation in cognitive capacity, with important implications for how cognitive, affective, and nociceptive processes are coordinated under high-demand conditions. Consistent with prior evidence that catastrophizing is associated with reduced IFG and may contribute to the link between negative cognitions and pain sensitivity [60], the present findings suggest that PTSS may similarly be associated with altered prefrontal engagement in chronic pain. Higher PTSS was associated with reduced dlPFC and IFGpo recruitment and increased coupling with threat-processing circuitry during task scans, suggesting the possibility that affective and trauma-related factors influence pathways involved in the regulation of both emotion and pain.

Chronic pain and PTSS are both factors individually known for reducing working memory performance. However, in our sample, neither PTSS nor chronic pain severity were correlated with working memory performance. Consistent with prior work in clinical populations, dlPFC activity in our sample did not predict working memory accuracy, highlighting that neural engagement can vary independently of task performance [26,53,76]. This pattern may also partly reflect the relatively restricted range of symptom severity and the modest sample size in the present cohort. Future studies with larger samples, broader clinical variability, and appropriate control groups will be needed to more fully characterize these associations; nonetheless, the current findings offer preliminary evidence for a potential neural pathway linking PTSS, cognitive control systems, and affective vulnerability in chronic pain. Taken together, the findings suggest that even subthreshold PTSS may influence neural processes known for their role in cognition, memory and regulation of pain and emotion.

Our findings show that greater functional connectivity between the left dlPFC and both the left dl/lPAG and vlPAG during task performance predicted reduced left dlPFC activation in the 3-back condition, as well as increased PTSS. Together, these associations suggest a pattern in which greater dlPFC-PAG connectivity co-occurs with reduced prefrontal engagement under high cognitive load in individuals with greater PTSS, highlighting the potential role of these regions in supporting the link between affective processing when cognitive load is high. Theories of working memory impairment in people with PTSD have proposed that heightened emotional distraction of PTSS contributes to cognitive dysfunction, likely due to reduced cognitive reserve [23,63,66,77]. The PAG is involved in threat detection [81], estimating threat probability [94,99], initiating defensive behaviours [22], and modulating responses to threats based on top-down contextualizing signals from the dlPFC [8,27,50,89]. Interactions between dlPFC and PAG have been implicated in shaping how executive functions such as working memory influence pain perception and threat response [87]. In addition, hyperconnectivity between these regions have also been reported in chronic back pain patients relative to healthy controls [52,102]. We previously demonstrated that dlPFC-PAG functional connectivity is associated with lower working memory and abnormalities in how the PAG processes threat signals in people with chronic pain, such that activations were abnormally elevated in individuals with lower working memory during conditions of positive expectation in a threat processing task [93]. Extending these findings, the current results suggest that increased dlPFC–PAG connectivity in the context of higher PTSS may reflect trauma-related alterations in how the brain coordinates cognitive and affective processes during working memory, providing a potential mechanism for individual variability in task-related neural engagement, and warrants further investigation.

In the present study, associations between PTSS and functional connectivity were observed for PAG subregions, but not for the amygdala or hippocampus. This supports the reports showing that the PAG plays a central role in descending pain modulatory pathways and the integration of cognitive, autonomic, and defensive responses, whereas the amygdala and hippocampus are more closely linked to contextual and evaluative aspects of affective processing rather than direct modulation of nociceptive signals [4,31,92]. Recent ultra-high-field (7 T) fMRI work from Wager and colleagues further supports this view, showing that the human PAG exhibits organized responses even during cognitive tasks in the absence of overt threat, with ventrolateral PAG engagement evident during working memory demands [28].

Increased functional connectivity between dlPFC and both PAG subregions during task correlated with total PTSS severity. More specifically, we found that dlPFC–dl/lPAG functional connectivity predicted arousal and reactivity symptoms, whereas dlPFC–vlPAG functional connectivity predicted avoidance symptoms. This aligns with previous reports of dl/lPAG and vlPAG involvement in active and passive coping, respectively, as well as previous reports of dl/lPAG driving the hyperarousal dimensions of PTSS [97] and the vlPAG driving the dissociative and avoidant dimension of PTSS [35]. While direct roles cannot be explicitly deduced, taken together with what is known about the functional differences of the regions, these findings highlight a pattern in which dlPFC–dl/lPAG connectivity is linked with arousal and reactivity symptoms, while dlPFC–vlPAG connectivity is linked with avoidance symptoms, potentially reflecting distinct roles of these circuits in different dimensions of PTSS. Importantly, this circuit-level distinction may help account for why PTSS is associated with increased affective, rather than sensory, pain, linking trauma-related hyperarousal and avoidance to the maintenance of heightened emotional responses to pain [32,40,90].

There are several limitations to consider when interpreting these findings. First, here we examine symptoms of post-traumatic stress independent of a PTSD diagnosis. Using a dimensional framework to evaluate PTSS allows symptoms to be measured with greater contextual sensitivity and captures differences in how symptoms manifest across individuals [15,55]. However, it does limit clinical generalizability to individuals with diagnosed PTSD. Relatedly, using the clinical cut off to identify people with high PTSS yielded only 14 participants, which reduces statistical power for between-group analyses. However, for most analyses when PTSS scores were treated as a continuous variable, the linear associations with PTSS were also significant. We also focused our interpretation on findings that were consistent across both analytic approaches. However, the small sample size should still be considered when interpreting comparisons. Finally, imaging the PAG can be challenging due to its small size and proximity to cerebrospinal fluid, which can reduce signal quality and increase susceptibility to noise. To account for these difficulties, we optimized a trade-off between voxel size and temporal resolution, using moderate multiband acceleration factors to maximize temporal sampling while preserving signal-to-noise ratio [17,95]. Importantly, the observed associations between PTSS clusters and connectivity of PAG subregions align with prior literature on dl/lPAG and vlPAG functional specialization [5,49] [93,95]. The biological relevance of these regions, as supported by prior research in humans and animals, aligns with our results. However, technical constraints related to imaging brainstem structures warrant caution, and further studies are needed to test the reproducibility of these findings and clarify the underlying mechanisms.

In conclusion, the findings from this study provide novel evidence that high PTSS, even at subthreshold levels for PTSD, are associated with alterations in neural mechanisms underlying working memory in individuals with chronic pain, highlighting early changes in brain circuitry that is associated with the affective components of chronic pain. The observed decrease in dlPFC activity, alongside increased dlPFC-PAG functional connectivity, highlight a pattern of neural interactions that may help in understanding how trauma-related symptoms are represented in working memory circuits, and that could reflect shift of cognitive resources even in the absence of measurable performance deficits. Further, the results provide evidence for the independent roles of PAG subregion connectivity with dlPFC, with dl/lPAG and vlPAG connectivity linked to arousal and reactivity symptoms and avoidance symptoms, respectively. Additionally, PTSS is associated with the affective dimension of pain, encompassing emotional responses such as fear, distress, and tension. This suggests heightened affective responses may interact with cognitive processes in chronic pain, potentially influencing top-down regulation of pain and emotion. Collectively, these results highlight neural patterns that may underlie working memory and affective vulnerabilities in individuals with chronic pain and PTSS, underscoring the importance of addressing both cognitive and emotional aspects of trauma in treatment approaches.

## Conflict of Interest Statement

The authors have no conflicts of interest to declare.

## Acknowledgements

This work was supported by the Natural Sciences and Research Engineering Council of Canada (NSERC) Discovery Grant [RGPIN-2016-05684], the Canada Research Chairs Program [CRC-2021-00164], the John R. Evans Leaders and Canada Innovation Funds (CFI-JELF) [CFI-35702], the Canadian Institute of Health Research (CIHR) Project Grant [PJT-168878], the Nova Scotia Health Authority (NSHA) Establishment Grant, the NSHA Fibromyalgia Research Grant, and the NSHA Anesthesia Research Fund.

## Supplemental Materials

**Supplementary Table 1.**
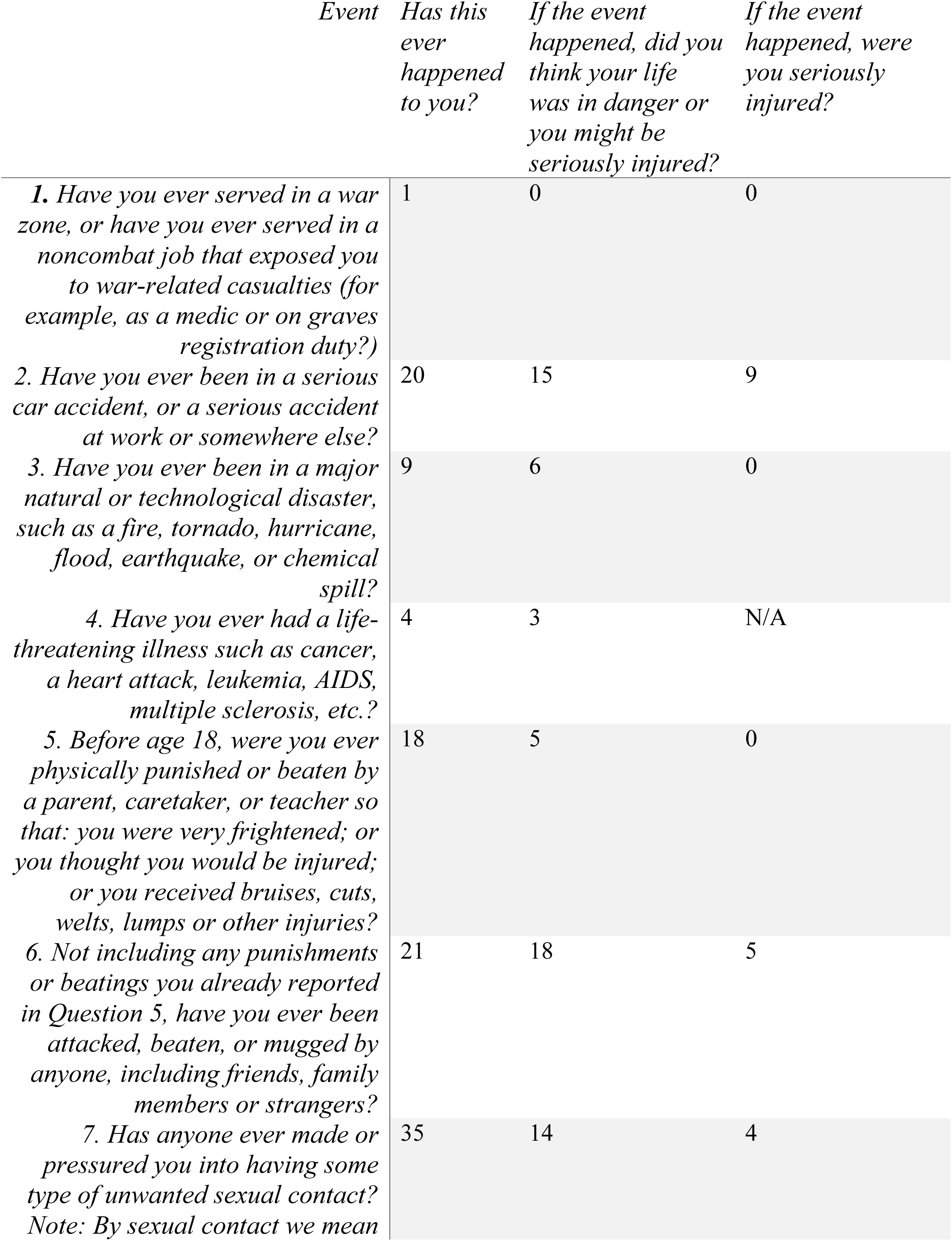
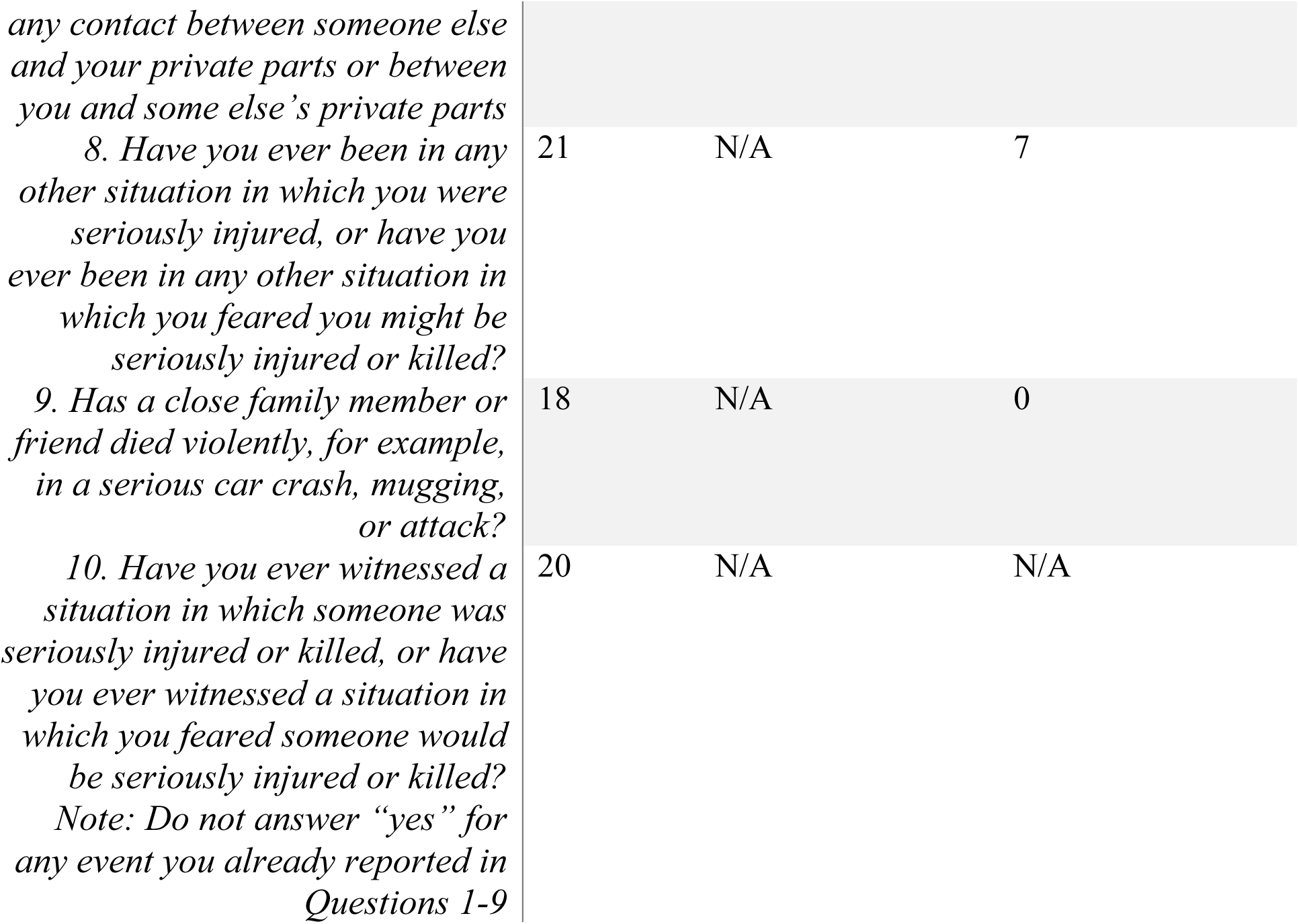
Number of participants who answered “yes” to each of the questions asked on the Brief Trauma Questionnaire. N/A= answer is considered not applicable according to the brief trauma questionnaire and so the question is not included in the survey.

## Notes

### Competing Interest Statement

The authors have declared no competing interest.

